# Higher eQTL power reveals signals that boost GWAS colocalization

**DOI:** 10.1101/2025.08.05.668745

**Authors:** Jonathan D Rosen, K Alaine Broadaway, Sarah M Brotman, Karen L Mohlke, Michael I Love

**Affiliations:** Department of Genetics, University of North Carolina, Chapel Hill, NC, 27599, USA; Department of Biostatistics, University of North Carolina, Chapel Hill, NC, 27599, USA

## Abstract

Expression quantitative trait locus (eQTL) studies in human cohorts typically detect at least one regulatory signal per gene, and have been proposed as a way to explain mechanisms of genetic liability for other traits, as discovered in genome-wide association studies (GWAS). In particular, eQTL signals may colocalize with GWAS signals, suggesting gene expression as a possible mediator. However, recent studies have noted colocalization occurs infrequently, even when expression is measured in biologically relevant tissues. Most eQTL studies to date include only hundreds of individuals, and are underpowered to discover distal regulatory signals explaining smaller fractions of gene expression variance. We integrate evidence from recent eQTL studies and demonstrate that limited statistical power due to sample size skews the detection of eQTL signals identified at various signal strengths. We estimate that a sample size of 500 detects <0.1 to 60% of eQTL for a range of signal strengths and that a sample size of 2,000 would detect 36.8% of all eQTL. We show that eQTL signals that can only be discovered in larger studies exhibit characteristics more similar to those of GWAS signals, including greater distance to the regulated gene and higher probability of loss intolerance. Finally, using results from recent eQTL studies and meta-analyses, we observe a large increase in detected colocalizations with GWAS signals compared to previous studies. These findings caution against overinterpreting the absence of colocalization in underpowered studies and provide guidance for designing future eQTL experiments, to improve power and complement perturbation-based approaches in characterizing gene-trait mechanisms.

## Main text

In the last 20 years, genome-wide association studies (GWAS) have successfully identified hundreds of thousands^1,2^ of associations between genetic variants and a diverse set of phenotypes. When GWAS signals (sets of linkage disequilibrium (LD)-linked associated variants) reside in coding regions and alter protein structure, a potential molecular mechanism underlying the association with a phenotype is apparent. However, the vast majority of GWAS signals are located in non-coding regions of the genome^3^, leading to one hypothesis that these associations are driven, at least in part, by the modulation of gene expression levels^4,5^. Similar in principle to GWAS, expression quantitative trait loci (eQTL) studies have been successful in identifying associations between genetic variants and gene expression in a variety of tissues and cell types^5–9^. Colocalization of GWAS and eQTL signals can then provide support for gene expression modulation as a mediator of GWAS association. Paradoxically, loci associated with GWAS traits often do not colocalize with eQTLs discovered in current studies, even when expression is assayed in a relevant tissue^10–14^, leaving many GWAS signals to be explained by other genetic mechanisms, future expression studies, or alternative assays. Several underlying reasons may contribute to this “colocalization gap”, including the measurement of gene expression in the wrong cell type, condition, or developmental stage, potential mediation of traits through RNA splicing instead of total gene expression, and low power for detection of eQTL.

Power has been a major contributor to the pace of GWAS signal discovery, and the same principle is expected to apply to eQTL signal discovery. Early GWAS, such as a 2007 WTCCC study, discovered 24 signals across 7 diseases with ∼17,000 subjects^15^. Recently, the GIANT consortium reported over 12,000 independent signals associated with height across 5.4 million subjects^16^, a sample size providing power to detect small effect sizes that was not feasible fifteen years prior. Despite debate about the explanatory value of such small effect sizes, GWAS loci have implicated genes and pathways with actionable therapeutic potential^17^. Several eQTL studies from easier-to-collect tissues have saturated the number of genes for which at least one eQTL signal can be found^10^ (“eGenes”). Brown et al^18^. noted over a decade ago that while eGene detection might reach saturation, the number of total eQTL signals had not. After discovering a primary signal associated with gene expression, additional signals can be discovered after conditioning on a variant representative of the primary signal, implemented in various stepwise statistical methods^6,19–22^. As eQTL studies grew larger, they also continued to find additional signals^5,23^. Thus, analogous to the pace of GWAS signal discovery, the full catalog of eQTL signals likely remains unknown at current power of eQTL studies.

Here, we integrate evidence from recent eQTL studies to demonstrate that limited statistical power from eQTL study sample size plays a key role in explaining the current GWAS-eQTL colocalization gap. We address the following three questions: 1) How much does increasing eQTL sample size increase the number of eQTL signals identified? 2) How do the characteristics of eQTL signals of various strengths compare to those of GWAS signals? 3) Would detection of additional eQTL from larger studies help to close the GWAS-eQTL colocalization gap? We answer these questions using data from four studies with sample sizes ranging from 103 to 4,703. With increases in sample size of recent eQTL studies and meta-analyses, we observe a large increase in detected colocalizations with GWAS signals, demonstrating the importance of eQTL study power in elucidating mechanisms of GWAS loci.

To systematically investigate eQTL detection power, we define a framework to quantify and interpret power across signals and studies. Signals with large effect sizes are easier to detect than small effects, and similarly, signals driven by common variants are easier to detect than those driven by rare variants^6^. Commonly used formulas for estimating detection power are functions of both effect size (beta) and minor allele frequency (MAF). The contributions of both beta and MAF can be represented by a single measure, Pearson’s correlation coefficient *r*, to quantify eQTL signal strength^24^ (Supplemental Material and Methods). Intuitively, larger beta and MAF values correspond to larger *r* values, *i.e.* stronger linear correlation between gene expression and genotype (Table S1). While beta is commonly used when discussing eQTL strength, *r* facilitates consistent comparisons across studies, and is interpretable as describing the variance in normalized expression explained by genotype (Figure S1).

We compare the distribution of *r* using primary eQTL signals detected in four studies: INTERVAL whole blood^7^ (n = 4,732), AdipoExpress subcutaneous adipose^6^ meta-analysis (n = 2,256), GTEx v10 subcutaneous adipose^5^ (n = 711), METSIM-S subcutaneous adipose^25^ (n = 420), and GTEx v10 kidney cortex^5^ (n = 103). The distribution of *r* for each study is shifted as expected according to the sample size of the study; *r* peaks at smaller values for larger studies, and large rightward shifts are observable between sample sizes in the hundreds, but the trend continues even with sample sizes in the thousands (Figure 1). We specifically compare AdipoExpress (larger n) and GTEx subcutaneous adipose (smaller n) due to their common tissue type and sample size difference. For common eGenes detected by both studies, the median *r* values are 0.133 and 0.140 for AdipoExpress and GTEx, respectively. In contrast, for eGenes detected by only one study, the median *r* values are 0.108 and 0.130, respectively (Figure S2). Thus, the distributional difference appears to be largely driven by the weaker signals detected in larger studies.

**Figure 1.**
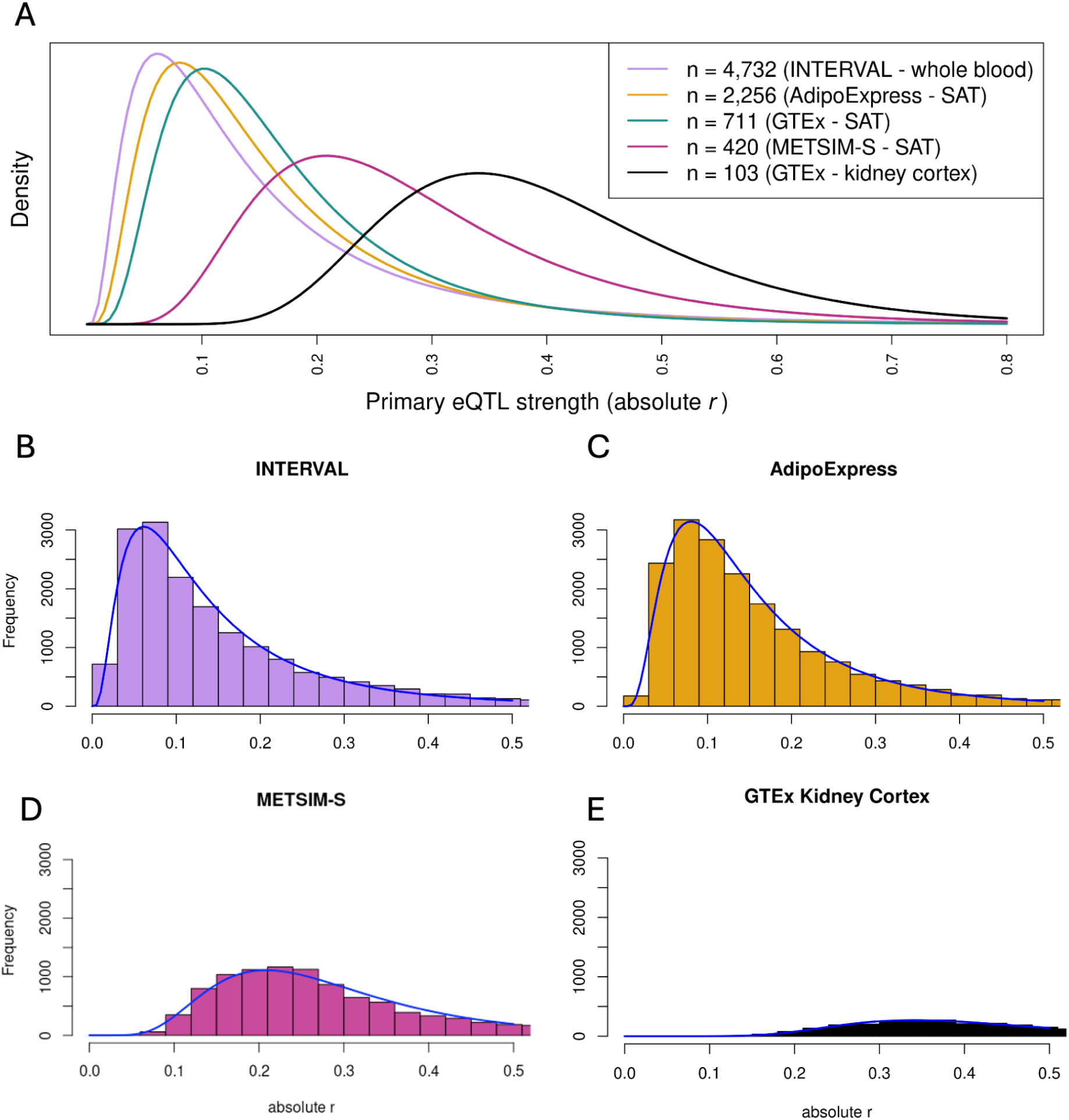
Distribution of observed primary eQTL signal strength. A) Fitted distribution of absolute value of r for primary eQTL signals in INTERVAL (n = 4,732), AdipoExpress (n = 2,256), GTEx subcutaneous adipose (n = 711), METSIM-S (n = 420), and GTEx kidney cortex (n = 103). B-E) Histograms of the empirical distributions of the absolute value of r for primary eQTL signals for four of the studies. Axes have the same scales across studies. The log-normal fitted distribution curve is plotted over the histogram to graphically display the quality of the fit. SAT - subcutaneous adipose tissue.

Using *r* values representative of ranges observed in the aforementioned studies, we can estimate the power to detect eQTLs at various sample sizes (Table 1, Supplemental Methods). Very strong signals (large *r*) can be detected with sample sizes in the hundreds (Table 2). At an *r* value of 0.2, corresponding to very strong eQTL signals (e.g. beta = 0.33, MAF = 0.25, 9th decile of detected eQTL signals in the INTERVAL study), the power at a sample size of 500 is still modest (<60% power). Even a sample size of 5,000 would have only ∼25% power for an *r* value of 0.05 (e.g. beta = 0.12, MAF = 0.10, a threshold that corresponds to the 37th quantile of detected eQTL signals in the INTERVAL study); more than 10,000 samples are required to detect an eQTL at 80% power with an *r* value of 0.05. Over 200,000 samples are required to reach 80% power at *r* = 0.01. This latter value represents ∼3% of the reported eQTL identified in the INTERVAL study. INTERVAL has only 1% power to detect QTLs at *r* = 0.01, suggesting that many eQTL signals remain to be detected. Together, these estimates show that many current eQTL studies are still underpowered to detect a large number of eQTL signals.

**Table 1.**
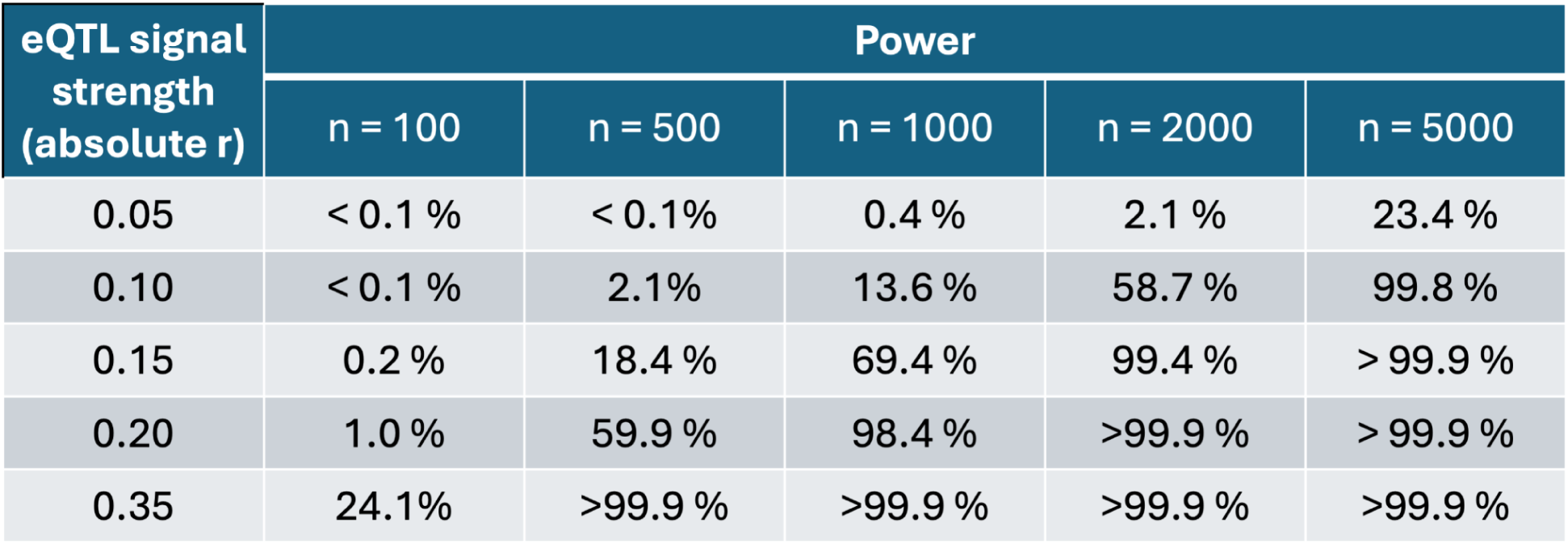
Estimated power to detect eQTL signals by signal strength and study sample size. Power estimates were calculated using a standard linear model assuming the response (gene expression) is standardized using a rank inverse normal transformation. n denotes eQTL study sample size. Refer to Table S2 for additional values.

| eQTL signal strength<br>(absolute r) | Power |  |  |  |  |
| --- | --- | --- | --- | --- | --- |
|  | n = 100 | n = 500 | n = 1000 | n = 2000 | n = 5000 |
| 0.05 | < 0.1 % | < 0.1% | 0.4 % | 2.1 % | 23.4 % |
| 0.10 | < 0.1 % | 2.1% | 13.6 % | 58.7 % | 99.8 % |
| 0.15 | 0.2 % | 18.4 % | 69.4 % | 99.4 % | > 99.9 % |
| 0.20 | 1.0 % | 59.9 % | 98.4 % | >99.9 % | > 99.9 % |
| 0.35 | 24.1% | >99.9 % | >99.9 % | >99.9 % | >99.9 % |

**Table 2.** Estimated sample size to achieve 80% power by eQTL signal strength. Power estimates were calculated using a standard linear model assuming the response (gene expression) is standardized using a rank inverse normal transformation.

| eQTL signal strength<br>(absolute r) | Sample size for 80%<br>power |
| --- | --- |
| 0.01 | 261,000 |
| 0.02 | 65,100 |
| 0.03 | 28,900 |
| 0.04 | 16,300 |
| 0.05 | 10,400 |
| 0.10 | 2,600 |
| 0.20 | 640 |

We next estimate the percentage of *all* eQTL signals detectable at a range of sample sizes. To do so, we fit a parametric distribution to the observed signal strength distribution of the INTERVAL study (Figure S3). We then use this fitted distribution and standard eQTL power formula to compute power for *all* signals, integrating across signal strength (Methods). At each signal strength, we can estimate the proportion of signals detected as the product of the proportion of signals (from the fitted distribution) and the probability of detecting such a signal (from power formula). Integrating this product over all signal strengths estimates the total proportion of all true signals detectable at a given sample size. At a sample size of 100, we predict that less than 5% of eQTL signals are detected, and at a sample size of 5,000, we estimate detection of only about half of eQTL signals (Table 3). As modeling signals based on detected signals biases the distribution towards stronger signals, we expect our current power estimates are overestimated. We note that power estimates at very small effect sizes are sensitive to small shifts in effect.

**Table 3.** Estimated percentage of total eQTLs detected at various study sample sizes. The estimated percentage is derived by integration of a convolution of power for a fixed effect size and the distribution of eQTL effect sizes. The distribution of power is based on a standard linear model, and the distribution of supposed “true” eQTL effect sizes is based on a log-normal fitted distribution using observed signal strength for all eQTL signals reported in INTERVAL.

| **n** | **Estimated percentage of eQTLs detected** |
| --- | --- |
| 100 | 3.6 % |
| 500 | 15.6 % |
| 1,000 | 25.0% |
| 2,000 | 36.8 % |
| 5,000 | 54.3 % |

To address the question of how the characteristics of eQTL signals of various signal strengths compare to characteristics of GWAS signals, we harmonize results across recent large studies. We examine results from the AdipoExpress and INTERVAL studies because both studies used conditional analysis to identify conditionally-distinct signals per gene, facilitating direct comparison of the entire sets of identified signals with conditionally distinct GWAS signals. Unsurprisingly, the larger INTERVAL study (n=4,732) detected 56,959 eQTL signals, substantially more than the 34,216 signals identified in AdipoExpress (n=2,256). These studies were conducted using different tissues, but the difference in the number of eQTL signals is likely driven primarily by sample size. We evaluate characteristics across deciles of eQTL signal strength (*r*), summarized in Table S1. For both studies, in line with previous reports^25–27^, we observe that lower strength eQTL signals tend to be located further from the transcription start site (TSS) of their associated gene (Figure 2).

**Figure 2.**
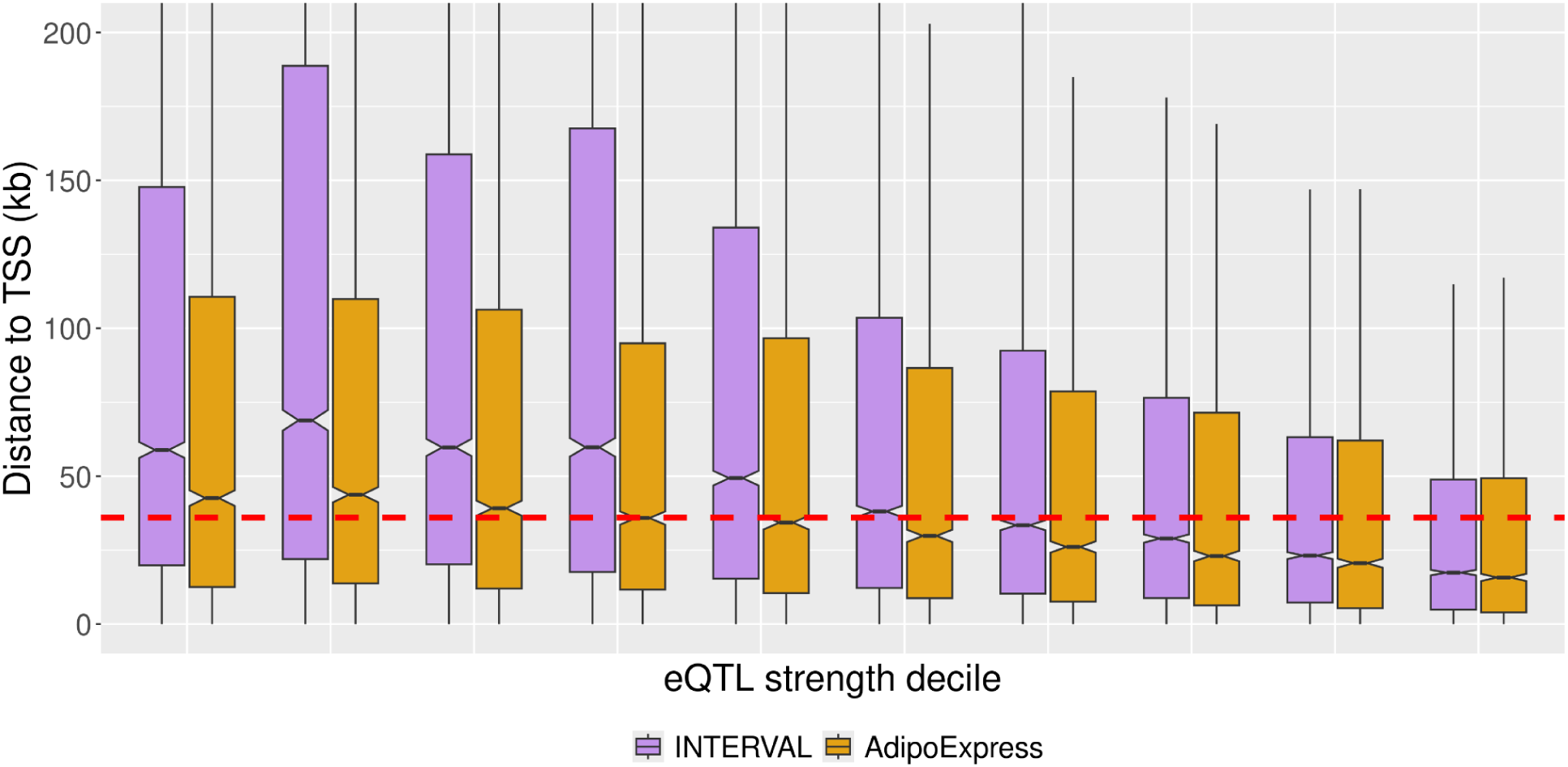
Distance to transcription start site (TSS) by signal strength. The x-axis shows deciles of absolute value of r, with weaker signal strength (smaller absolute r) on the left, and stronger signal strength on the right. The dashed line at 36 kb represents the median distance of a GWAS variant to the nearest gene TSS as reported by Mostafavi et al.^12^ The points represent the median TSS for the corresponding bin, and the lines span the 25th and 75th percentiles. Purple - INTERVAL, yellow - AdipoExpress. The plot is truncated at 200 kb for clarity.

We next compare the distances of eQTL lead variants to TSS observed for these eQTL studies to those reported in GWAS. The median variant-to-TSS distances of the strongest decile of eQTL signals in the AdipoExpress and INTERVAL studies are 16 and 17 kb, respectively. The median distances for the weakest decile are approximately three-fold higher, 43 and 59 kb, respectively. Recently, Mostafavi et al^12^. reported the median distance of a GWAS lead variant to the closest gene TSS was 36 kb. In addition, they reported a median distance of 13 kb from eQTL variants to the closest gene TSS. We surmise that this short distance reported for eQTL to gene is due both to the use of the closest gene TSS (as opposed to associated gene TSS) and the limited sample size of GTEx v8. With a more comprehensive picture of the distribution of eQTL signals from higher powered studies, we find that approximately 52% of INTERVAL signals and 45% of AdipoExpress signals are located at signal-to-gene distances equal to or greater than the reported median GWAS-to-gene distances (*e.g.* 36 kb).

We examine if similar trends can be observed with respect to four other characteristics of GWAS and eQTL signals^12^: the percent of lead variants in promoter regions, percent of lead variants in distal enhancers, percent of associated genes with high probability of loss intolerance (pLI), and percent of associated genes that encode transcription factors (TF). Mostafavi et al. reported^12^ that GWAS signals differ systematically from eQTL signals in that GWAS signals are less likely to overlap promoters, more likely to overlap distal enhancers, and are more often adjacent to high pLI genes and genes that encode TFs. In our analyses, weaker (smaller *r*) eQTL signals are more GWAS-like than stronger ones: weaker signals are less likely to overlap promoter regions, more likely to overlap distal enhancer regions^26^, and more likely to be associated with high pLI genes and genes encoding TFs (Figure S4). eQTL signals that can only be discovered in larger studies exhibit characteristics more similar to those of GWAS signals.

To address whether increased eQTL detection power can help close the GWAS-eQTL colocalization gap, we performed colocalization analyses (Supplemental Methods) using GWAS signals from five continuous traits (total cholesterol, triglycerides, waist-hip ratio adjusted for body mass index, and high and low density lipoprotein-C) and two dichotomous disease phenotypes (type 2 diabetes and coronary artery disease) with subcutaneous adipose eQTL from AdipoExpress^6^ (n=2,256, 34,216 eQTL signals) and one of its smaller component studies, METSIM-S (n=420, 13,392 eQTL signals). On average, the number of colocalizing GWAS signals approximately doubles when using the meta-analysis eQTL data compared to the smaller dataset (Figure 3). For instance, a chromosome 19 GWAS signal for waist-hip ratio adjusted for BMI colocalizes with an eQTL signal for *NFIX* with a small *r* value (0.045; first decile) detectable only in AdipoExpress (Figure S5). Notably, the increase in colocalizations is not solely driven by stronger eQTLs: across deciles of eQTL signal strength, the proportion of colocalizing signals remains relatively stable in both AdipoExpress and INTERVAL (Figure 4). These findings show that higher-powered eQTL studies can uncover additional GWAS-eQTL colocalizations, including those involving weaker eQTL signals.

**Figure 3.**
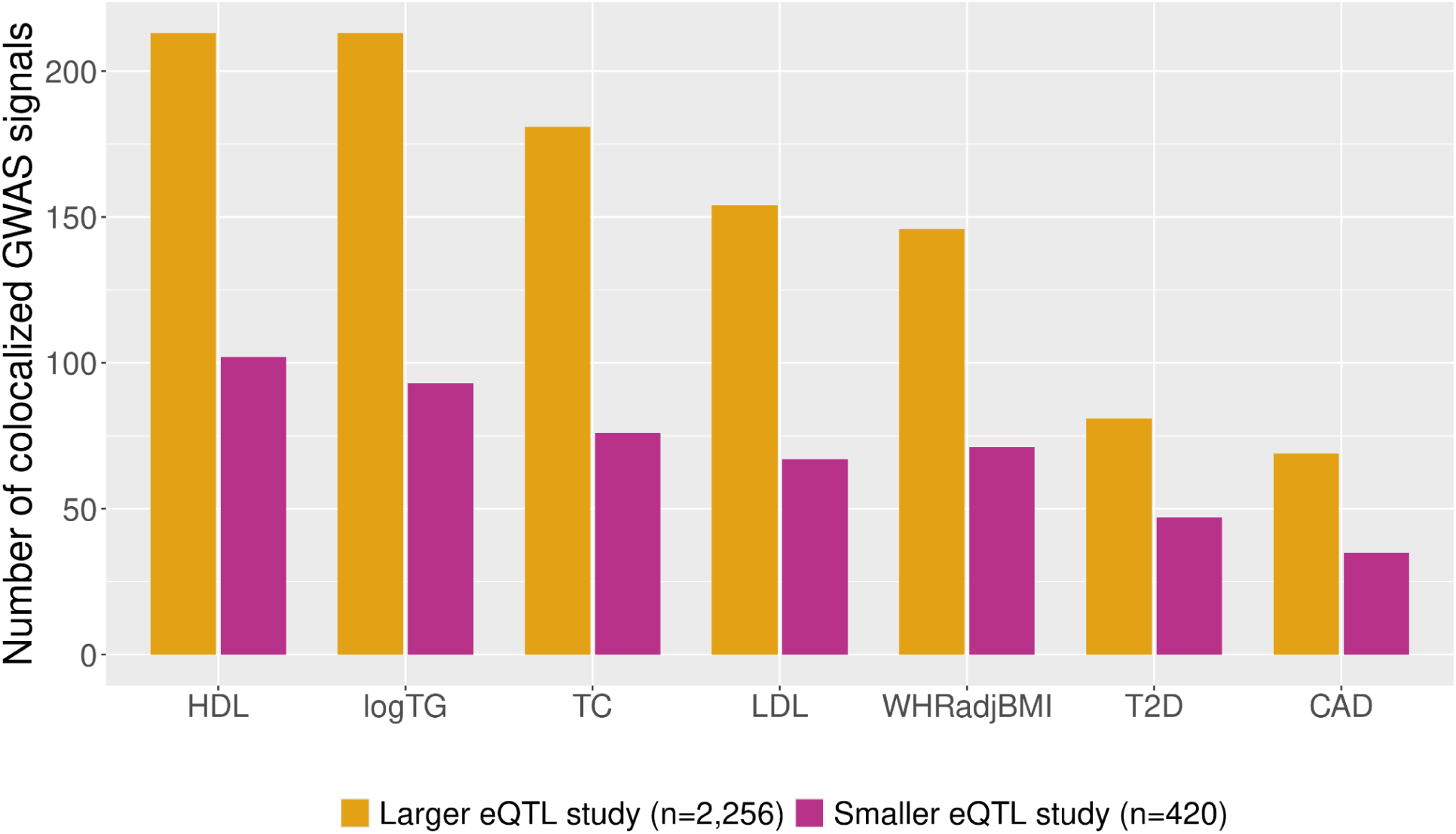
Comparison of the number of colocalized GWAS signals detected using parallel methods by eQTL study size. The height of each bar corresponds to the number of GWAS signals colocalized (PPH4≥0.5) with at least one eQTL in each study (yellow - AdipoExpress, pink - METSIM-S). HDL: High-density lipoprotein cholesterol, logTG: log triglycerides, TC: total cholesterol, LDL: Low-density lipoprotein cholesterol, WHRadjBMI: waist-hip ratio adjusted for body mass index, T2D: type 2 diabetes, CAD: coronary artery disease. Mean fold change per trait = 2.1 (range 1.7 - 2.4).

**Figure 4.**
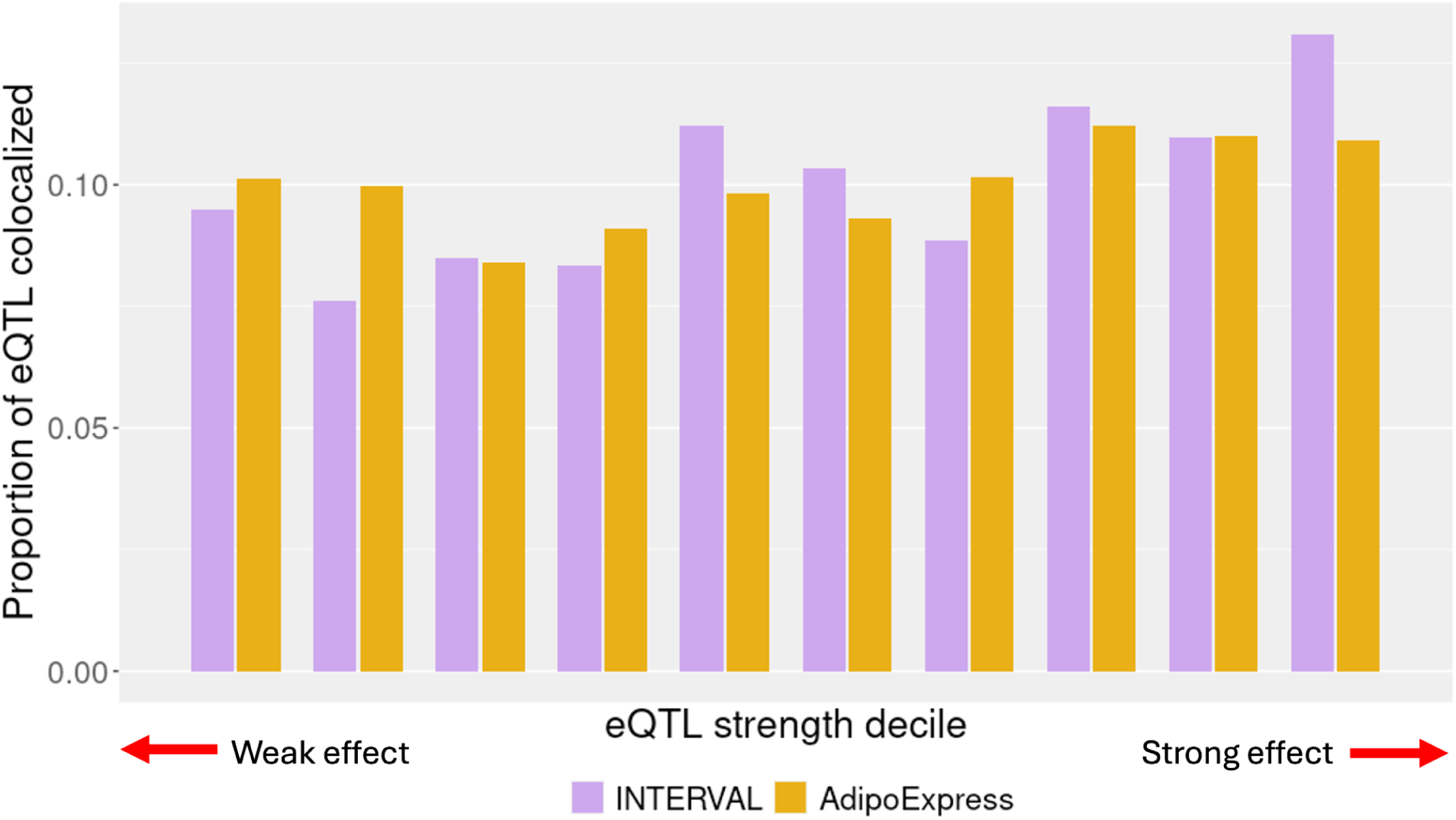
Proportion of eQTL signals that colocalize with a GWAS trait by signal strength. The height of each bar corresponds to the proportion of eQTL signals that colocalize with at least one GWAS signal across all traits examined. Purple - INTERVAL, yellow - AdipoExpress.

Furthermore, the increase in colocalizations is not apparently driven by a higher yield of strong eQTL signals in the larger study. Within deciles of eQTL signal strength, we compute the proportion of eQTL signals that colocalize with at least one GWAS signal. In both AdipoExpress and INTERVAL, the proportions of colocalizing eQTL signals are relatively consistent across eQTL signal strength (Figure 4; ranges defined in Table 4). Given that statistical methods used to define colocalization are sensitive to eQTL signal significance, which in turn is driven by effect size, it is possible that even more colocalization with weaker eQTL signals would be observed in better powered studies.

**Table 4.** Range of values of *r* for indicated deciles. For the AdipoExpress and INTERVAL studies, the lower and upper values of *r* for each decile are shown. Intervals are closed on the right and open on the left, i.e. (a, b].

| **Decile** | **INTERVAL** |  | **AdipoExpress** |  |
| --- | --- | --- | --- | --- |
|  | Lower (*r*) | Upper (*r*) | Lower (*r*) | Upper (*r*) |
| 1 | 0 | 0.021 | 0 | 0.048 |
| 2 | 0.021 | 0.032 | 0.048 | 0.061 |
| 3 | 0.032 | 0.042 | 0.061 | 0.073 |
| 4 | 0.042 | 0.054 | 0.073 | 0.087 |
| 5 | 0.054 | 0.067 | 0.087 | 0.102 |
| 6 | 0.067 | 0.084 | 0.102 | 0.120 |
| 7 | 0.084 | 0.108 | 0.120 | 0.146 |
| 8 | 0.108 | 0.115 | 0.146 | 0.184 |
| 9 | 0.115 | 0.224 | 0.184 | 0.258 |
| 10 | 0.224 | 1 | 0.258 | 1 |

Together, these analyses show that discovering additional eQTL signals, including weak and distal ones, further informs mechanisms for GWAS signals. Here, we have aggregated effect sizes from recent eQTL studies to show that the power for discovery of many eQTLs remains low at common eQTL study sample sizes. Recent large eQTL studies have reported that the majority of genes harbor multiple eQTL signals^6,7,23^ and therefore, a single detected eQTL signal per gene does not saturate eQTL discovery. We found that weaker signals only detected in larger studies exhibit characteristics (distance to TSS, and properties of the associated gene) that are more similar to GWAS signals than the stronger eQTL signals found in previous studies. Across the distribution of eQTL signal strength, the signals detected only in larger studies continue to provide additional colocalization with GWAS signals. In a recent whole blood eQTL study using TOPMed by Orchard et al^23^., the authors note that the majority of colocalizing GWAS signals (67%) do so with at least one secondary eQTL signal. They also demonstrate far more colocalizations comparing TOPMed eQTL signals to GTEx, observing that GWAS signals colocalizing with TOPMed but not GTEx tended to do so with weaker cis-eQTL signals, consistent with our analysis (Figure S2). Weaker eQTL signals identified in bulk tissue through conditional analysis may colocalize with GWAS loci even when stronger eQTL signals do not. This scenario may reflect limitations of statistical colocalization methods or the fact that bulk eQTLs aggregate signals across heterogeneous cell types. Regulatory variants may modulate gene expression in a relevant cell type that affects downstream trait or disease risk, while variants belonging to another signal may modulate expression in a cell type that has no such effect.

Our findings help avoid misinterpreting limited evidence of GWAS-eQTL colocalization from underpowered studies as indicating the lack of expression mechanisms, and help to inform design of future studies. While eQTL studies with sample sizes in the thousands take time, planning, money, and effort, meta-analysis can combine results across multiple studies, increasing effective sample size beyond what might be feasible in a single study. QTL studies that deeply explore expression across single cells or across multiple timepoints or conditions are invaluable for uncovering novel regulatory contexts and informing cell-type and context-specific predictive models; such studies are nevertheless subject to the same logic about sample size and discovery potential as for bulk eQTL and GWAS signal colocalization.

Continued identification of eQTL signals with more moderate effect sizes has additional benefits. As in GWAS, even loci with small effects on expression can provide information about gene pathways and potential therapeutic targets. Identifying additional eQTL signals increases the pool of GWAS-eQTL signal pairs for colocalization testing, and approaches for gene prioritization leveraging colocalization have been shown to provide additional specificity compared to other methods such as TWAS^28–31^. Colocalizing pairs can inform the direction and magnitude of the gene-to-trait effect in disease-relevant tissues^6,9,32–34^.

In addition to more powerful eQTL studies, further discovery is needed of context-specific eQTLs in which genetic regulation is only observable in one or a few cell types, tissues, environments, or conditions, as these would be missed in well-powered studies of mismatched contexts^35–39^. Splicing and other molecular QTL also have been shown to explain a substantial portion of GWAS signals^40,41^. The use of complementary assays such as CRISPRi/a^42,43^ will also help to decipher mechanisms driving additional GWAS associations and provide additional evidence for or against eQTL-signal-based models.

Human eQTL studies contribute unique information among complementary assays. They offer the chance to observe the impact of genetic variation on gene regulation in the natural state: embedded in interacting cell types, tissues and environments, compounded by the potential impact of genetic regulation across development or cell differentiation. Such studies are vital to fully understanding many disease etiologies and ultimately developing therapies.

## Declaration of interests

The authors declare no competing interests.

## Acknowledgments

We acknowledge support from National Institutes of Health grants UM1 HG012003, R01 DK072193, R01 DK093757, UM1 DK126185, R01 DK132775, T32 HL129982, and F31 HL154730. The Genotype-Tissue Expression (GTEx) Project was supported by the Common Fund of the Office of the Director of the National Institutes of Health and by NCI, NHGRI, NHLBI, NIDA, NIMH, and NINDS. We thank Jason Stein and Doug Phanstiel for helpful discussions.

## Web resources

INTERVAL: https://www.intervalrna.org.uk/downloads/

AdipoExpress: https://zenodo.org/records/13845120

GTEx: https://www.gtexportal.org/home/downloads/adult-gtex/qtl

TF genes: http://humantfs.ccbr.utoronto.ca

ENCODE cCRE: https://screen.encodeproject.org/

pLI: https://gnomad.broadinstitute.org/downloads

LiftOver: https://genome-store.ucsc.edu/

## Data and code availability

AdipoExpress: https://zenodo.org/records/13845120

INTERVAL: https://zenodo.org/records/10354433

## Supplemental Figures

**Figure S1.**
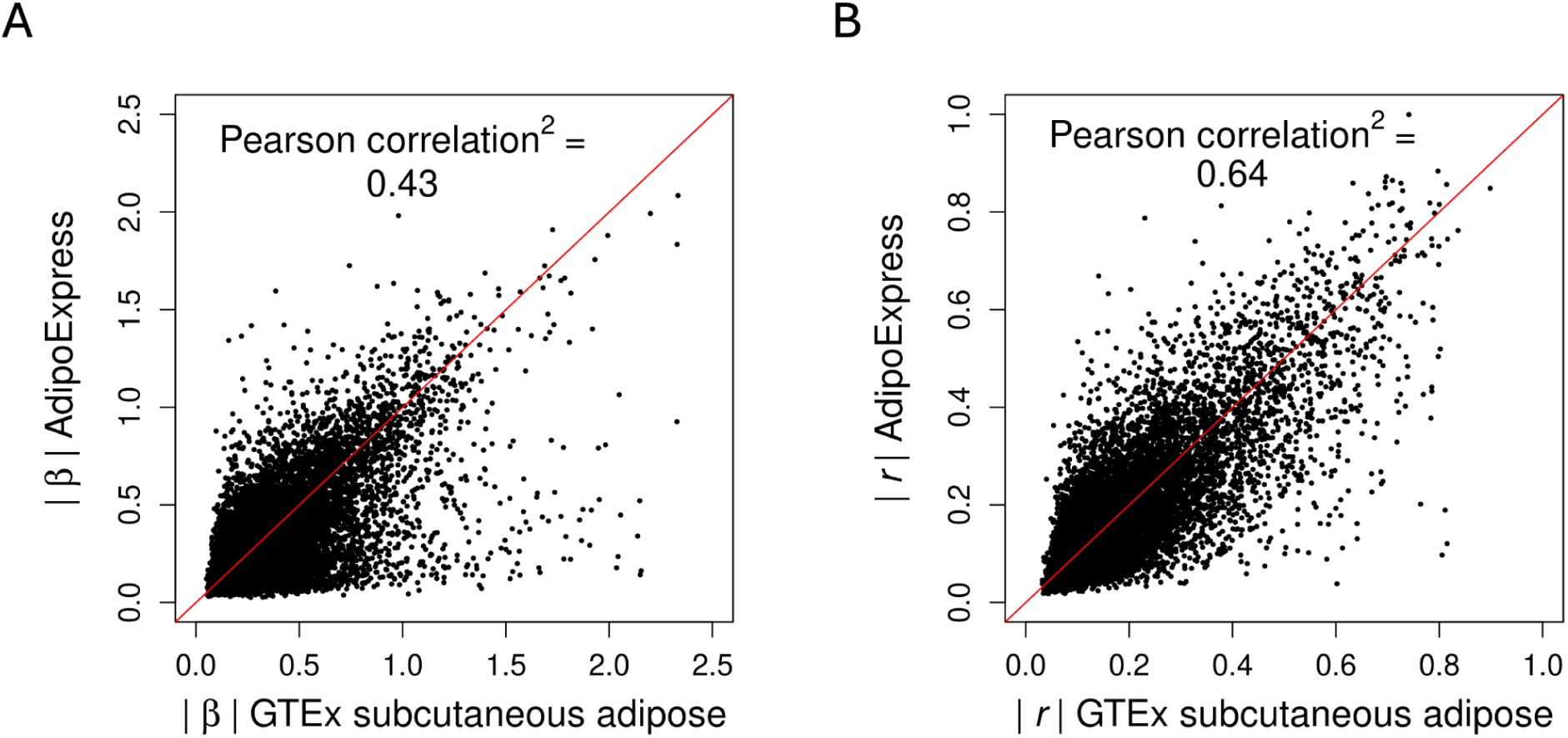
Concordance of effect sizes between AdipoExpress and GTEx subcutaneous adipose. A) Comparison of the absolute values of the (A) beta coefficients and (B) r value for primary eQTL signals for overlapping genes between GTEx v10 subcutaneous adipose (x-axis) and AdipoExpress (y-axis).

**Figure S2.**
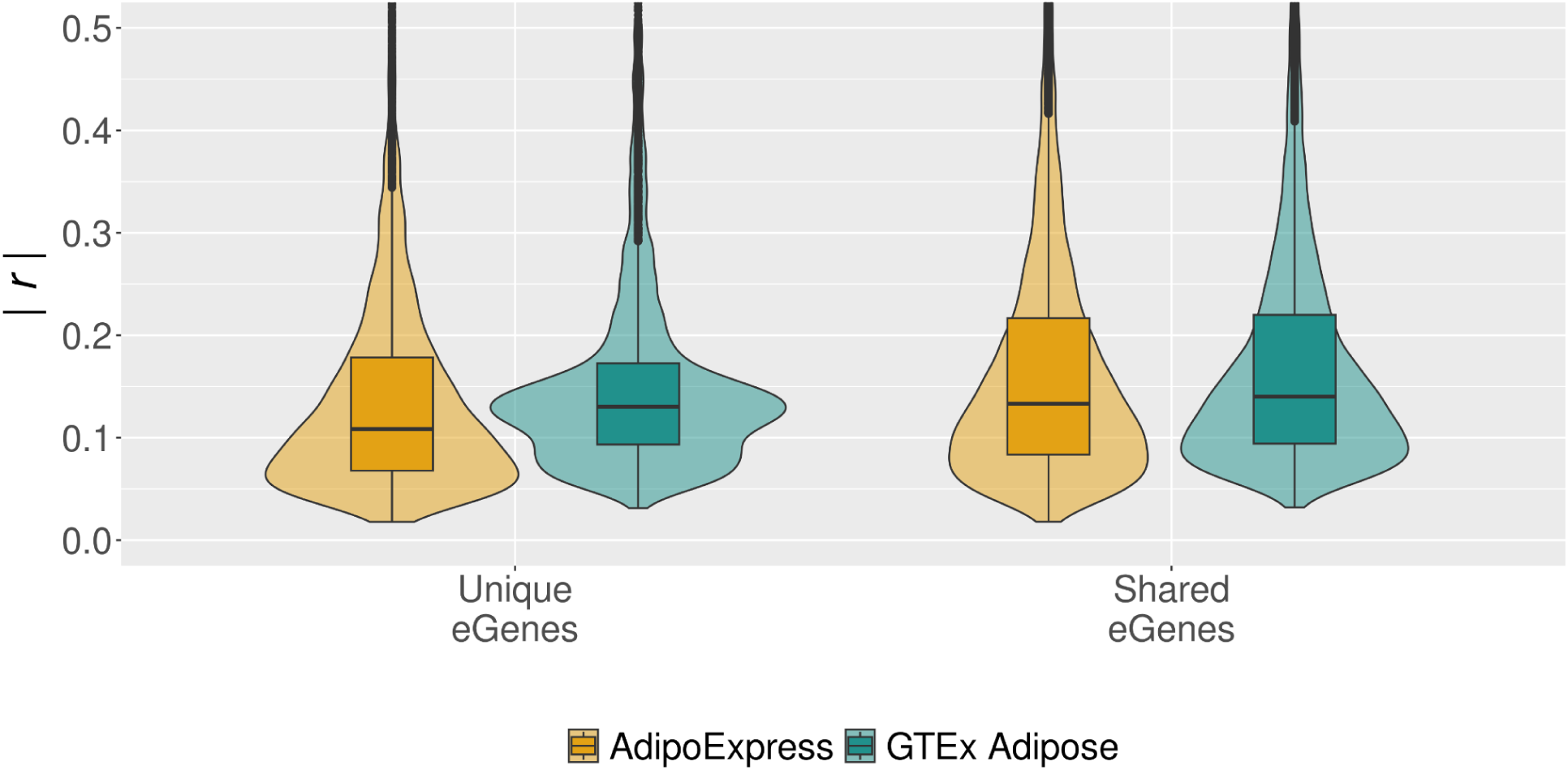
Comparison of effect size between AdipoExpress and GTEx subcutaneous adipose. Absolute value of *r* (y-axis) for primary signals in the set of overlapping eGenes (right) and non-overlapping genes (left) for both AdipoExpress and GTEx v10 subcutaneous adipose. The top, middle, and bottom of the boxplot inside each violin represents the 75th, 50th and 25th percentiles of the values, respectively.

**Figure S3.**
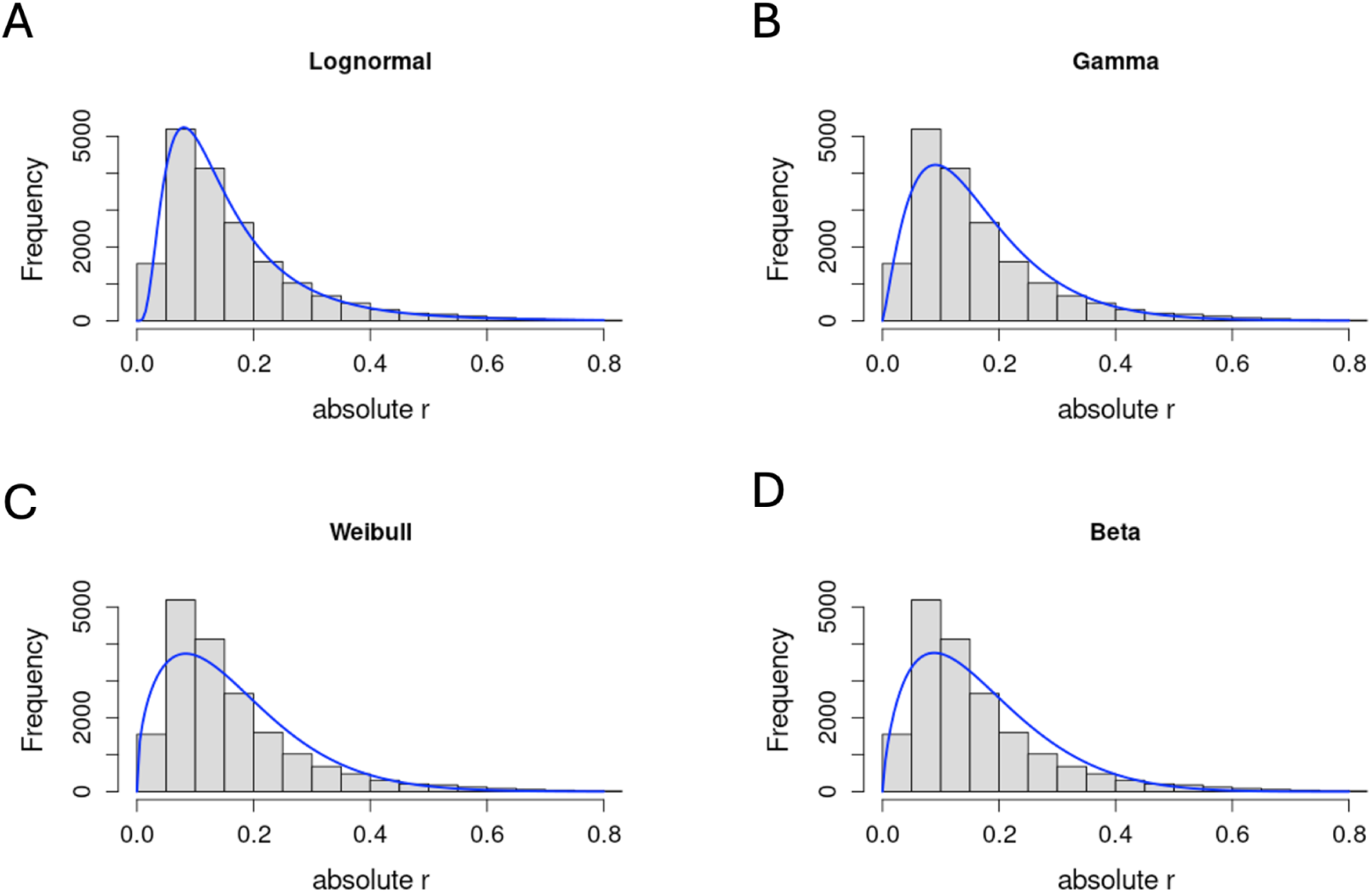
Fitting parametric distributions to an empirical distribution of eQTL effect sizes. A-D) Histograms of observed absolute r for primary signals in AdipoExpress with fitted distributions overlaid (blue lines). Distributions are lognormal (A), gamma (B), Weibull (C), and beta (D) and were fitted using the “fit” function in R.

**Figure S4.**
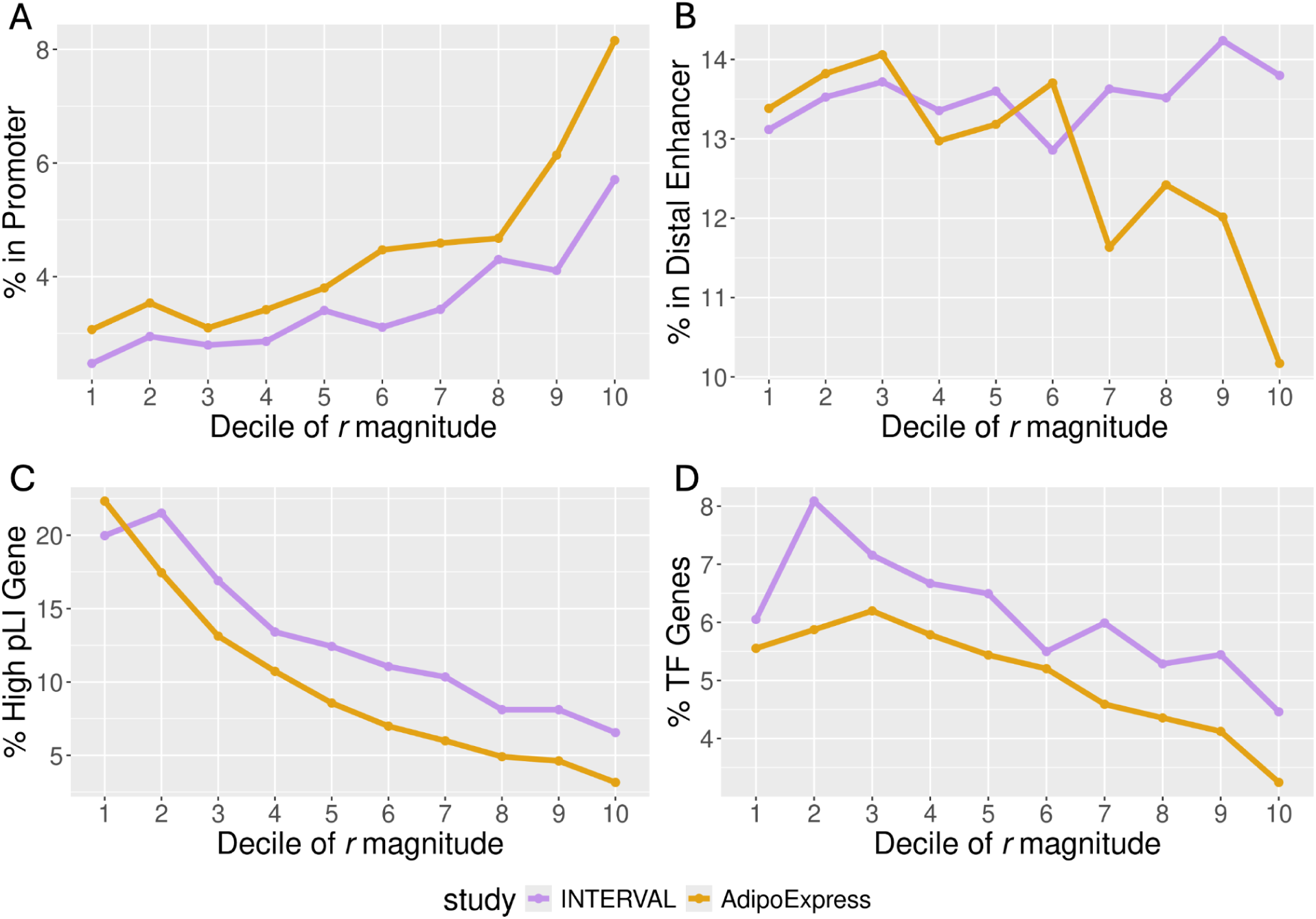
Trends in characteristics of eQTL signals by signal strength. Shown over the decile of r magnitude (x-axis, increasing magnitude moving from left to right) are the following: (A) percentage of signals per decile located within an ENCODE annotated promoter region or (B) distal enhancer region and (C) percentage of eGenes associated with the eQTL signal that are classified as high pLI genes or (D) transcription factors.

**Figure S5.**
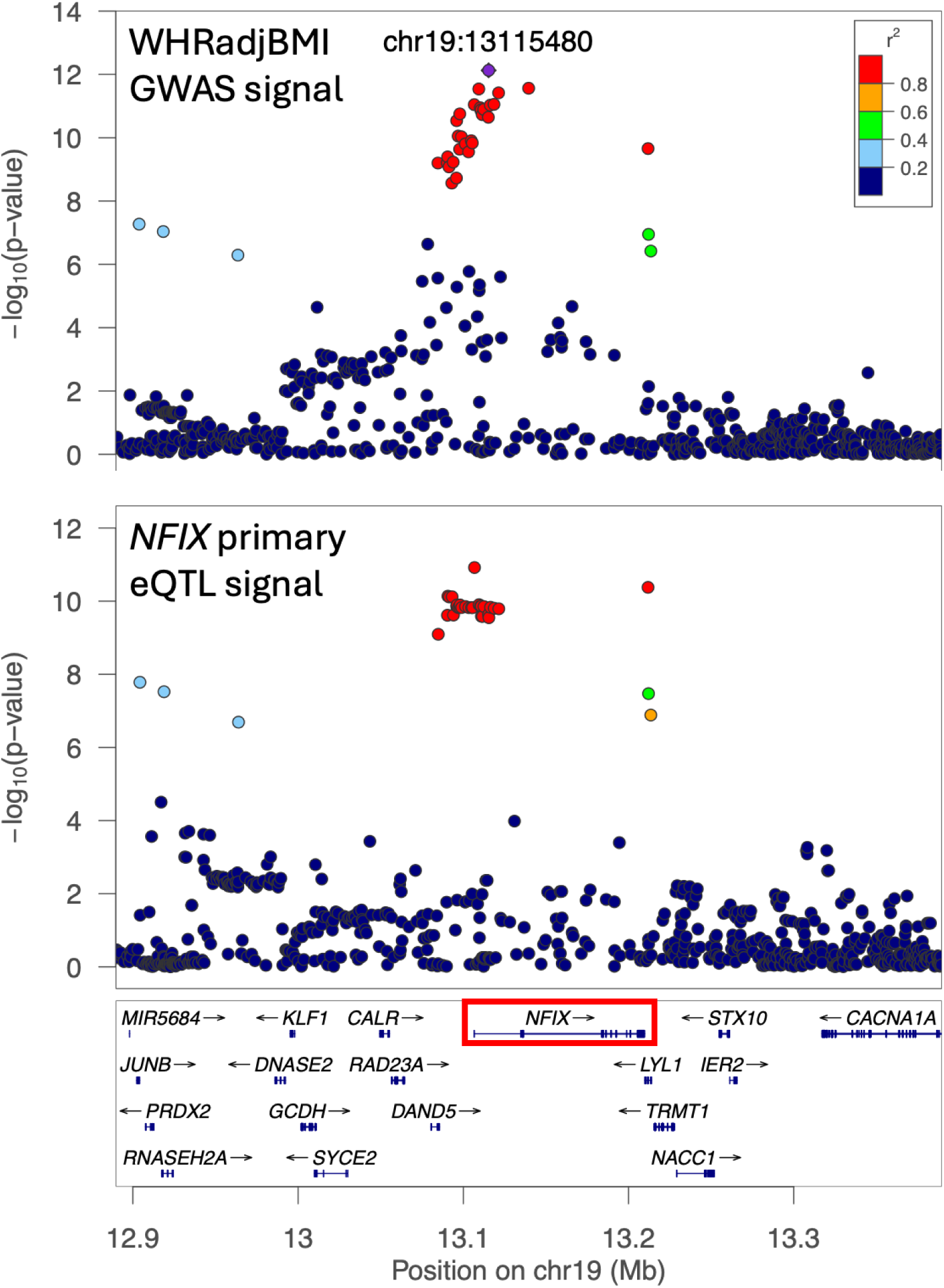
LocusZoom plot of colocalized GWAS-eQTL signals. Top: GWAS signal for waist-hip ratio adjusted for BMI (ref); Bottom: primary eQTL signal for *NFIX*. The eQTL signal was identified in the larger AdipoExpress study but not in METSIM-S. The lead variant for the eQTL signal has *r* = 0.046 (beta = −0.21, MAF = 0.025).

## Supplemental Material and Methods

### Use of Pearson correlation for computing power

Power to detect an eQTL signal is a function of both the linear coefficient describing the direction and magnitude of effect of genotype on gene expression (β) and the minor allele frequency (MAF) of the variant^44^. This dependence of power on two parameters limits comparisons of power estimates because one of the two parameters is typically held constant to observe the effect of the other on power. However, having a single metric to describe eQTL signal strength that accounts for the contribution of both β and MAF is desirable.

We assume a standard linear model

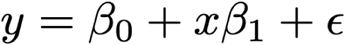

where y denotes gene expression, X denotes genotype (0, 1, 2 copies of allele), and ε is an error term assumed to be normally distributed. Many eQTL analysis pipelines standardize gene expression to fit such a model by performing a rank-based inverse normal transformation on scaled RNA counts such that y is normally distributed with mean 0 and variance 1. Under Hardy-Weinberg equilibrium, the standard deviation of X can be expressed as

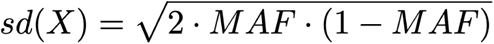

Under this framework, the sample Pearson correlation (r) between genotype (X) and gene expression (y) can be expressed as

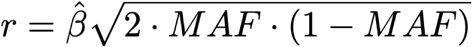

since

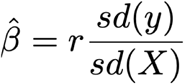

and sd(y) = 1. This strategy was employed by Vosa et al^24^ when estimating power in the eQTLGen study.

We use the R package powr to perform power calculations based on r. Given our ability to model *Pr*(|r|), the distribution of the absolute value of r, we can then estimate the percentage of eQTL detected at various sample sizes using the convolution:

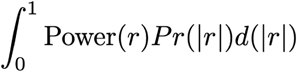

### Comparing signals across GTEx and AdipoExpress

The majority of the eGenes exclusive to GTEx were not tested in AdipoExpress, so it is expected the median *r* is close to that for the common eGenes. The AdipoExpress meta-analysis required genes to exceed an expression threshold in at least two studies to be included. One potential reason for lack of expression in studies other than GTEx is the different location of the fat depot. The majority of eGenes exclusive to AdipoExpress that have significantly lower *r* values were tested in GTEx but not found to be significant.

### Overlap with promoter or distal enhancer region

From the SCREEN registry of cCREs v3^45^, all human cCREs (hg38) were downloaded as a single .bed file and lifted over to hg19 using the UCSC LiftOver tool. An eQTL signal was considered to overlap with a promoter region if any nucleotide of the lead variant was located within a region annotated as “PLS” or “PLS,CTCF-bound”. An eQTL signal was considered to overlap with a distal enhancer region if any nucleotide of the lead variant was located within a region annotated as “dELS” or “dELS,CTCF-bound”. Percent overlap by decile was determined by first ranking all eQTL signals by magnitude of the absolute value of r and then partitioning the entire set of eQTL signals into ten equally sized bins by rank. The number of overlapping eQTL signals divided by all eQTL signals per bin was then determined. Since the bins are equally sized, the percent overlap can be used to compare absolute counts across bins as well.

### Percent high pLI or transcription factor genes

The file “gnomad.v2.1.1.lof_metrics.by_gene.txt.bgz” was downloaded from the Broad Institute website (https://gnomad.broadinstitute.org/downloads) and Ensembl IDs for genes with pLI scores ≥ 0.9 were extracted into a list of high pLI genes. eQTL signals were linked to their associated gene, which was cross-referenced with the list of high pLI genes to determine percentages per decile. The TF gene overlap was determined analogously after extracting Ensembl ID from the corresponding list.

### Colocalization

For each trait, we used PLINK (v.1.90b3) to calculate the LD *r*^2^ between all conditionally distinct GWAS lead variants and conditionally distinct adipose eQTL lead variants within 500 kb of each other using 40,000 unrelated UK Biobank (UKBB) participants as the LD reference panel^46^. If the LD *r*^2^ was ≥0.5, we tested GWAS–eQTL pairs for colocalization using coloc (v.5.1.0.1, coloc.abf, default settings). We considered GWAS–eQTL signal pairs colocalized if the coloc PP4 was ≥0.5. For the METSIM-S comparisons, a separate colocalization analysis was not performed. Rather, we computed the LD (PLINK v.1.90b3) between all pairwise combinations of eQTLs in the smaller study with those in the meta-analysis for each eGene. A GWAS signal was considered colocalized with a METSIM-S eQTL if LD *r*^2^ was ≥0.5 with a colocalized meta-analysis eQTL for the same gene.

## Notes

https://zenodo.org/records/13845120

### Competing Interest Statement

The authors have declared no competing interest.

